# Let there be nightlights: the ecological role of bioluminescence in a Costa Rican mushroom

**DOI:** 10.1101/2023.01.10.523474

**Authors:** Catharine A Adams, Marion Donald, Christin Anderson, Efraín Escudero, Susanne Sourell, Sven Landrein, Carolina Seas, Greg Mueller, Priscilla Chaverri

## Abstract

Both marine and terrestrial organisms produce light enzymatically in a process called bioluminescence. Though the ecological role of light production is known for some species, such as fireflies and bacteria, the ecological role in mushroom-producing fungi remains relatively unexplored, particularly in Central America. Here, we discovered a bioluminescent mushroom in the high-elevation oak forests of Costa Rica. We developed mushroom models with Light-Emitting Diodes (LEDs) producing various colors: green, blue, red, and yellow. Over three consecutive evenings after sunset, we applied Tanglefoot invertebrate-trapping glue to both mushroom models and actual mushrooms, and collected the traps before sunrise, then identified the trapped invertebrates to Order. We found green LED traps attracted more invertebrates than non-lit control traps, suggesting that light functions to attract invertebrates. The majority of invertebrates attracted to the green lights were Dipteran flies, who would be capable of dispersing fungal spores. The higher-intensity green LEDs attracted more total invertebrates than the dimmer mushrooms, but the results were not significant. Though we predicted that the invertebrate assemblages attracted to green lights would be similar to the invertebrate assemblage attracted to actual mushrooms, the results were not significant. Similarly, the red, blue, and yellow LEDs attracted fewer invertebrates than the green LEDs, but the differences in community composition were not statistically significant. Our findings corroborate other similar studies in tropical regions that found bioluminescent mushrooms may attract invertebrates.

## Introduction

A diverse array of organisms biochemically produce light in a chemical reaction known as bioluminescence. Enzymatic light production evolved well over 40 times across the tree of life (Fleiss and Sarkisyan 2019)). Though the majority of bioluminescent organisms are found in the ocean (Haddock et al. 2010; Davis et al. 2016), bioluminescence can also be found among land-dwelling microbes (Wilson and Hastings 1998), insects (Meyer-Rochow 2007; Jeng 2019), and fungi (Stevani et al. 2013; Tsarkova et al. 2016). Bioluminescence is typically produced by the oxidation of a light-emitting substrate, generally called the luciferin. The reaction is catalyzed by an enzyme, usually termed a luciferase, to produce an oxyluciferin and a visible photon (Mager and Tu 1995).

The ecological role of bioluminescence varies greatly depending on the organism in question. Some organisms such as fireflies produce light to signal to potential mates (Lewis and Cratsley 2008), others like dinoflagellates signal to scare potential predators (Latz and Rohr 2005), and microbes like bacteria use light for quorum sensing to regulate cell density (Bassler et al. 1997). Among fungi, three separate lineages of mushrooms produce their own light: the *Armillaria* lineage (Mihail 2013), the *Omphalotus* lineage (Bondar et al. 2011), and the mycenoid lineage (Desjardin et al. 2008). In total, over 100 species of mushroom produce light, and mushrooms from each lineage share the same type of luciferin and luciferase (Oliveira et al. 2012). However, the ecological role of bioluminescence in mushrooms remains relatively unexplored.

Three main hypotheses attempt to explain the ecological role of bioluminescence in mushrooms. The first hypothesis states that bioluminescence is merely a byproduct of metabolism (Sivinski 1981; Weitz 2004). Lignin degradation has been linked to bioluminescence, and may function to detoxify peroxides formed during the process (Bermudes et al. 1991).The second hypothesis posits that the light serves in a protective role, to deter harmful fungivores or attract predators of fungivores (Sivinski 1981). The third hypothesis postulates that light serves to attract potential spore dispersers. One older study in Florida, USA (Sivinski 1998) and a recent study in Brazil found support for hypothesis 3 (Oliveira et al. 2015) but these results have yet to be replicated in Central America.

Here, we designed an experiment to explore the ecological role of mushroom bioluminescence in the high-elevation oak forest of Costa Rica. We describe what may be the first instance of a bioluminescent mushroom in the forest surrounding the Cuericí Biological Station. We then designed an experiment to test the following hypotheses 1) model ‘mushrooms’ with green Light-Emitting Diodes (LEDs) attract more insects than their dark controls; 2) model mushrooms with green LEDs attract similar assemblages of insects as actual mushrooms that emit a similar wavelength of light; 3) the higher-intensity green LED attracts a greater number of total insects than actual mushrooms; and 4) red, blue, and yellow LEDs attract fewer insects, and different assemblages of insects as green LEDs, since they have no nearby naturally-occurring mushroom equivalent.

## Methods

### Study Location

The study was conducted during the wet season in 2015 in the tropical cloud forest at Cuericí Biological Station (9.555000°, −83.667778°), in the Cartago province of Costa Rica. At an elevation of 3000 m, the forest canopy is predominantly oak, *Quercus costaricensis*. The mid-story is composed of plants in the Araliaceae and Lauraceae families as well as palms, bamboo, and tree ferns. The understory consists of ferns, vascular epiphytes, bryophytes, and plants in the Araceae, Compositae, and Solanaceae families.

### Mushroom specimen collection

One hour after sunset, we set out to obtain bioluminescent mushrooms. Walking along an established trail, we would periodically turn off all light sources, wait several minutes for our eyes to adjust to the absence of human-made light, and then scan visually for bioluminescence. A *Mycena*-like mushroom species was found at several sites along the trail, growing out of dead oak. Several specimens of different sizes were brought back to the field station for taxonomic description.

### Mushroom model fabrication

Clear polycarbonate was cut into strips 4 cm wide by 11 cm long. Each piece was bent lengthwise into a table-like shape such that each table ‘leg’ was approximately 3 cm tall, and the width of the top of the table was 5 cm across. Next, a hole was drilled into the center of each table to allow for an LED to rest snugly on the top of the table surface, and electrical tape was used to fasten a 3V Lithium battery to the cathode and anode of the LED for constituent light production (**Figure 1**). To control for the effect of LED color, mushroom models were made with LEDs that emit either green light, blue light, yellow light, red light, or no light (control).

**Figure 1:**
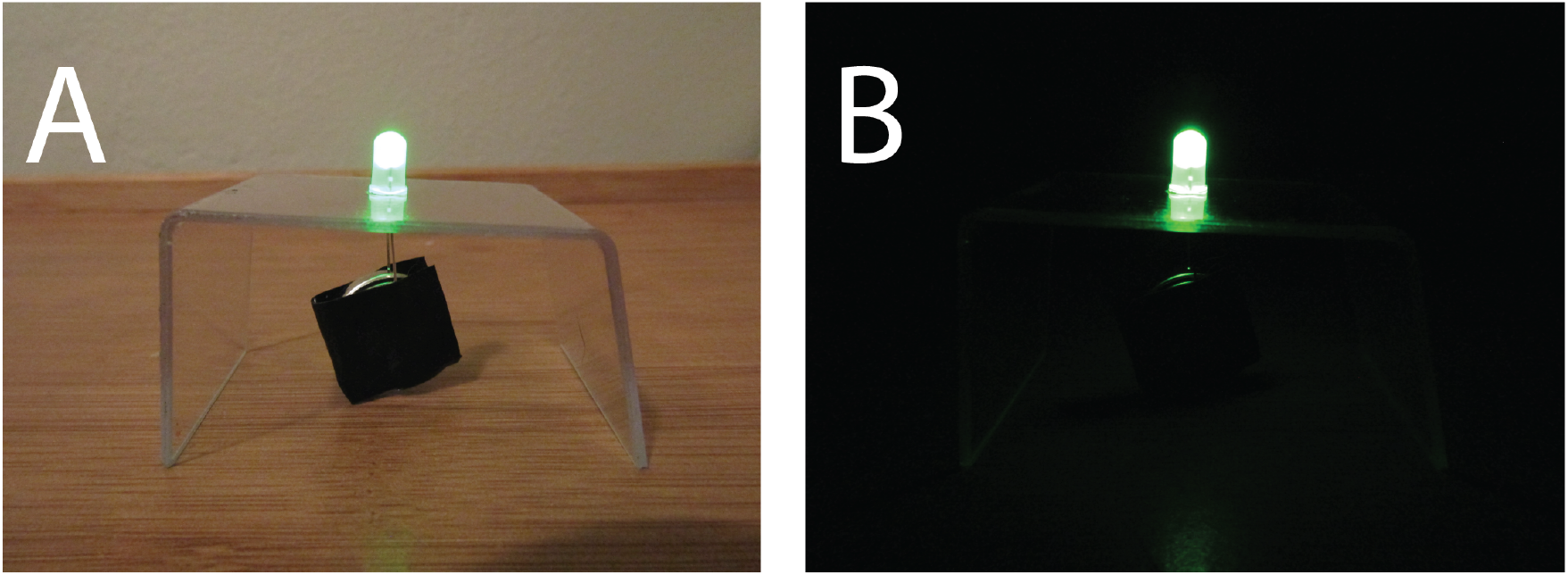
Mushroom model A) with overhead lighting, and B) without overhead lighting.

### Experimental design

Two to three hours after sunset, one color of each mushroom model, as well as a non-lit control, were placed near a patch of mushrooms. Each mushroom model was situated 15-20 cm apart, and the top of each table was coated with a thick layer of Tanglefoot insect barrier©, as was 1 nearby mushroom (**Figure 2**). Mushrooms and mushroom models were collected at least 30 minutes before sunrise. The experiment was replicated three times on three separate nights, and at two nearby sites each night, for a total of six replicates.

**Figure 2:**
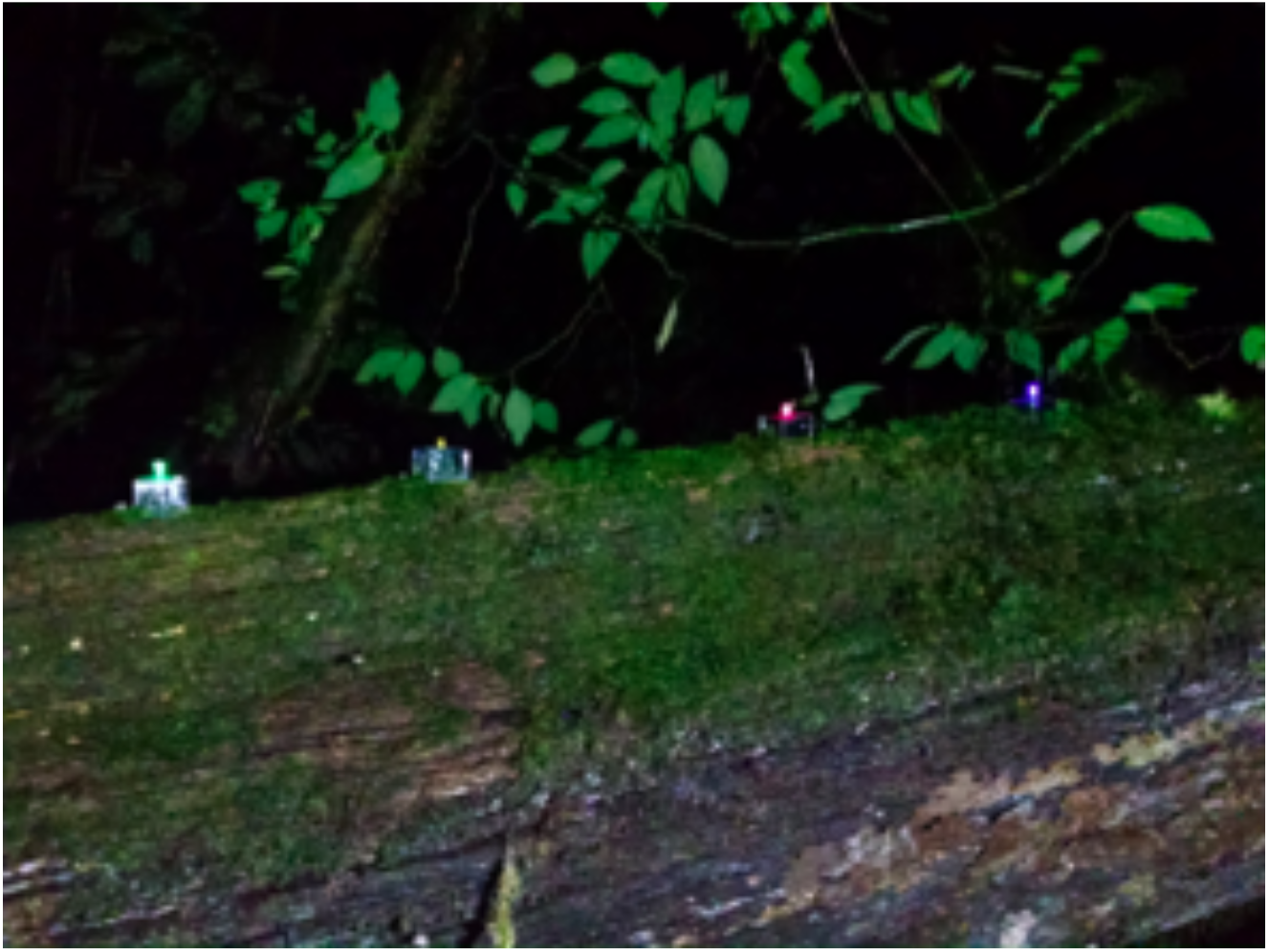
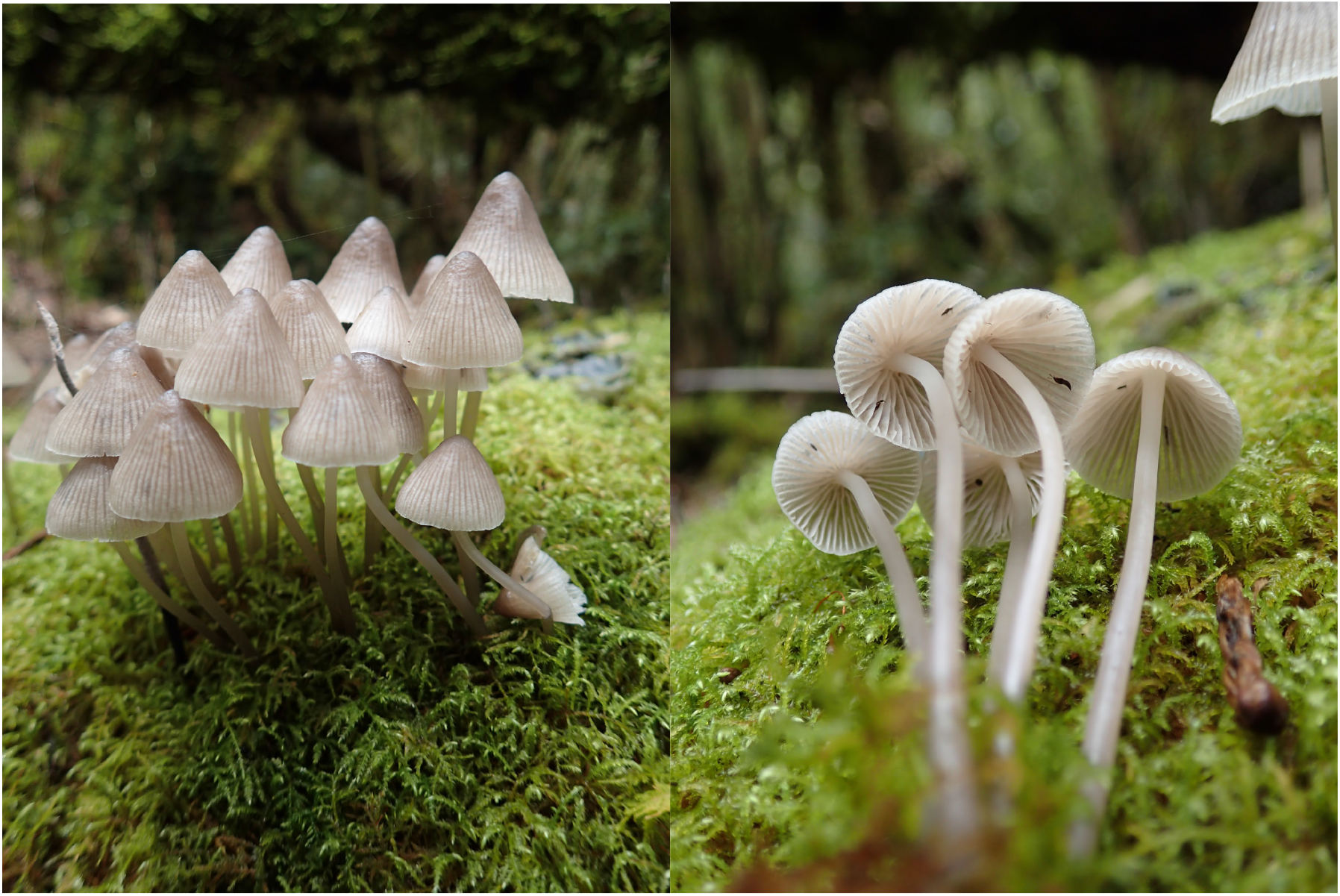
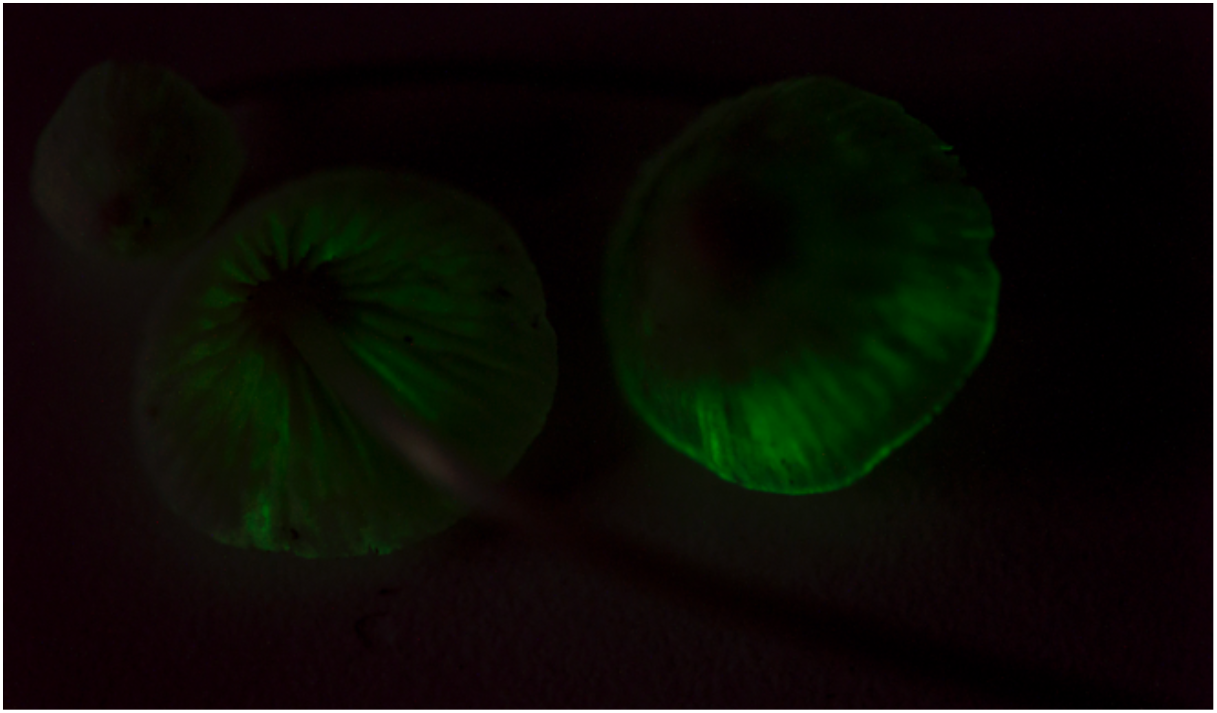
An example of experimental set-up in the field. Figure 2A: *Mycena sypherii* nom. prov., top view. B: *Mycena sypherii* nom prov., pileus underside. Photos credit Susanne Sourell C: Bioluminescent pileus underside (left) and top (right). Photo credit Efraín Escudero.

### Invertebrate identification

Invertebrates were removed from the mushroom models or mushroom with forceps as carefully as possible, examined with dissecting microscopes, counted, and identified to Order. When the invertebrates could not be identified, they were classified as “Unidentified.”

### Analysis

To understand the influence of trap type on invertebrate community composition, we first visualized log-transformed invertebrate communities using Bray-Curtis distances with NMDS. We then used PERMANOVA to determine if trap type predicts insect community composition. As our control lightless traps attracted no invertebrates, this trap type was excluded from the analyses. These analyses all relied on the R-package *vegan* (Oksanen et al. 2019) v.2.5.6. and plots were visualized in *ggplot2* (Wickham 2016) within the computing environment R v.3.6.1 (R Core Team 2019).

## Results

### Mushroom description

The pileus is gray-brown, becoming translucent-white at margin; broadly conical, glabrous, 8 to 15 mm in diameter. The flesh is fragile with a mild odor and taste. The stipe is hollow, watery; 38 to 51 mm long, 0.5-1.5 mm wide; equal, white at apex, the same color as the pileus. The lamellae are broadly distant, off-white, and <1 mm deep. The lamelluae are 1 to 2-seried. There is scant white basal mycelium. The lamellae and pileus margin glow greenish-white in the dark.

The hymenium has four-spored basidia. The spores are amyloid, subglobose to broadly ellipsoid, with prominent Hilar appendages. The pileus trama is strongly dextroid. The hyphae possess clamp connections. The mushroom will hitherto be referred to as *Mycena sypherii* nom. prov.

Because the data were not normally distributed, to determine whether green traps attracted more insects than control traps, we performed a Mann-Whitney U test. The difference was significant (p = .02).

We next asked if the brighter green traps attracted more insects than mushrooms. Again, because the data were not normally distributed, we performed a Mann-Whitney U test, and found that the difference was not statistically significant (p =.23).

To determine whether trap type explains insect community composition, we used Permanova (‘adonis’ in vegan) (Oksanen et al 2015). We did not detect a statistically significant difference in community composition across the trap types (R^2^ = .3, p = .385). We did not find statistically significant support that green LEDs attract similar assemblages of insects as mushrooms. Nor did we find support that blue, red, or yellow LEDs attract different community assemblages from green LEDs and mushrooms.

## Discussion

Here we find evidence supporting the notion that mushroom bioluminescence functions at least partially to attract insects. We found support for our main hypothesis, Hypothesis 1: the green LED mushroom models attracted more insects than the dark controls, which did not trap a single insect (Mann-Whitney U test, p = .02). Hypothesis 2 was not supported: Green LEDs attracted a somewhat similar assemblage of insects as actual mushrooms (Figure 5), but the results were not statistically significant (R^2^ = .3, p = .39). For Hypothesis 3, the higher-intensity green LEDs attracted more total insects than their dimmer mushroom counterparts (Figure 3), but the results were not significant (Mann-Whitney U test, p = .23). Finally, hypothesis 4 was not entirely supported: the red, blue, and yellow LEDs attracted fewer insects than the green LEDs (Figure 3), but the differences in community composition were not statistically significant (Figure 5).

**Figure 3:**
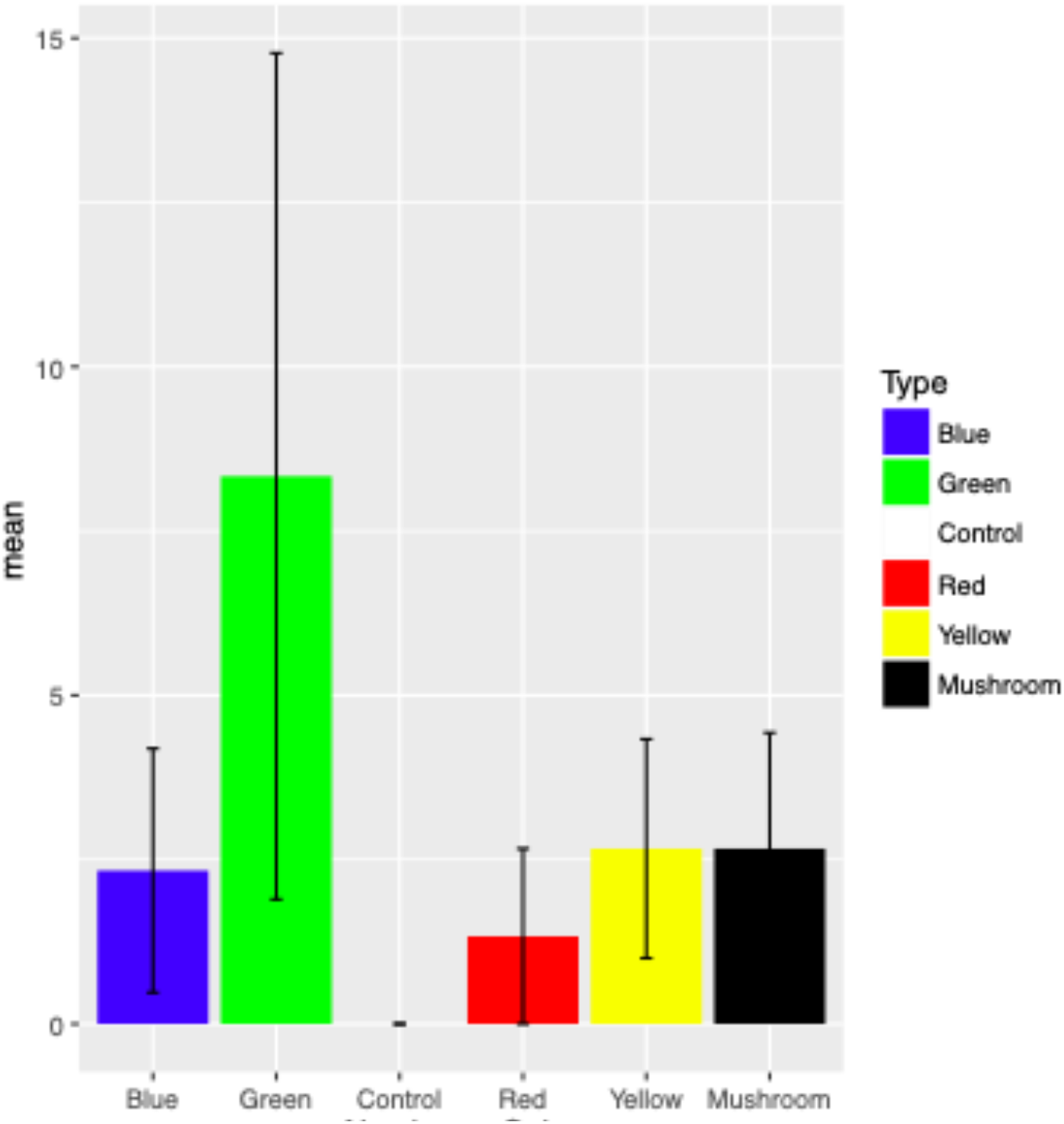
Average number of total insects caught across all three nights at Cuericí. Error bars are standard deviations.

**Figure 4:**
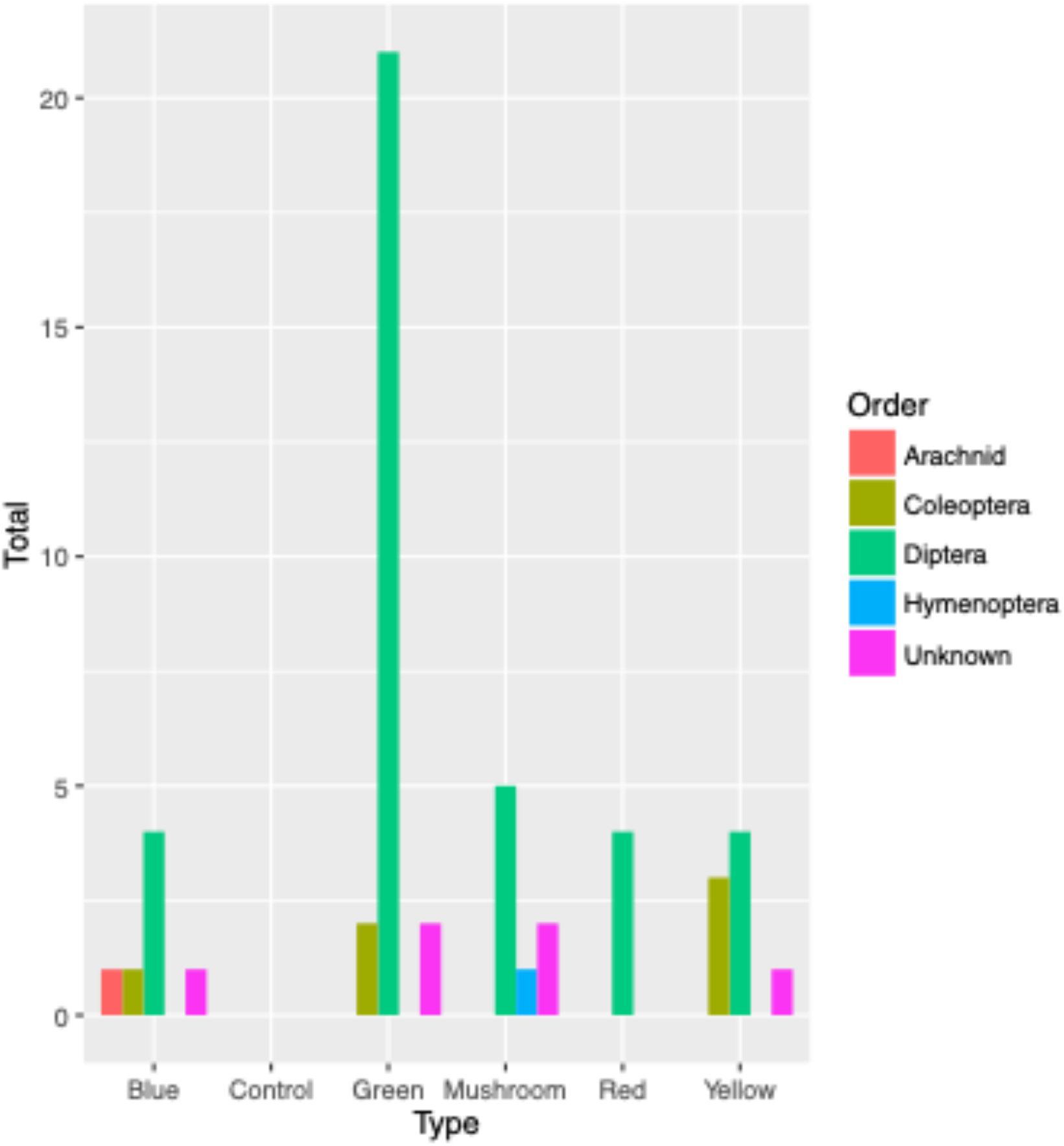
Total number of each Order of insect found on each light and/or actual mushroom at Cuericí.

**Figure 5:**
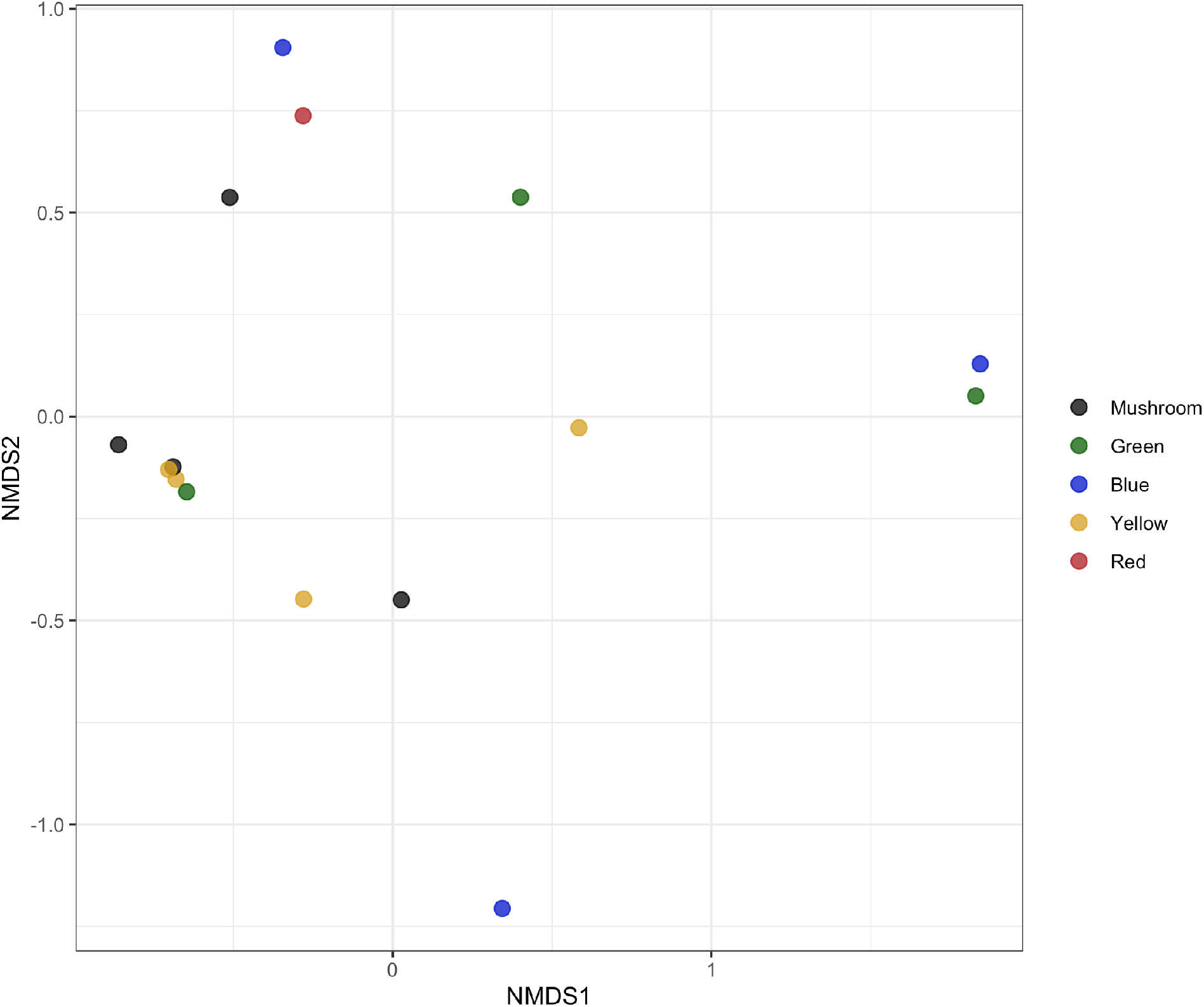
NDMS on log-transformed abundance data with Bray-Curtis dissimilarity matrix. Stress = 0.08468, k = 2. Points are jittered 0.05 in height and width to decrease complete overlap and visualize all communities.

Support for Hypothesis 1 is evidenced by the complete lack of insects on the unlit controls, compared to both mushrooms and the green LED traps. These findings corroborate two previous studies with comparable experimental design. In Florida, USA, Sivinski found that both glowing *Mycena* sp., mycelium and bioluminescent fruit bodies of *Dictyopanus pusillus* attracted more insects than their respective unlit controls (Sivinski 1981). Research in Brazil with *Neonothopanus gardneri* discovered that unlit mushroom model controls attracted a small number of Diptera and some Hymenopteran insects, but no Coleoptera or Hemiptera, unlike lit (green) mushroom models which attracted insects of all four Orders (Oliveira et al. 2015). Together, these studies support the hypothesis that mushroom bioluminescence functions to attract insects.

However, the ecological role of fungal bioluminescence, or at least its relative importance, may be region-specific. For example, an experimental study in temperate Australia found no significant differences in insect attraction between lit and unlit treatments (Weinstein et al. 2016). Their findings may be partially because the authors used pieces of bioluminescent *Omphalotus nidiformis* instead of entire mushrooms.

Another possible explanation is that insect dispersal may be more important in dense tropical forests, where wind flow can be restricted near to the ground (Oliveira et al. 2015). In a similar vein, many North American bioluminescent mushrooms tend to be dimmer than their tropical counterparts (Sivinski 1981), further suggesting light may play a larger role in tropical forests.

Both green LEDs and actual mushrooms attracted more Diptera than any other insect Order (Figure 4). Being winged, flies and other Dipteran insects are particularly capable of dispersing spores. However, the sample sizes were not large enough to conclude whether the attracted insect assemblages were significantly similar. This experiment should be repeated with more LED traps at more sites, over more nights. Future research should also test whether insects that associate with bioluminescent mushrooms truly move spores away from the parent mushroom.

This is the first experiment, to our knowledge, to compare insect attraction of bioluminescent mushrooms and green LEDs to lights that are blue, red, and yellow. The red LEDs only attracted Diptera species (Figure 4), while blue attracted more different insect Orders than both green LEDs and actual mushrooms: Arachnid, Coleoptera, Diptera, and Unknown. The yellow LEDs attracted Diptera, Coleoptera, and Unknown insects, much like the green LEDs. Regardless, none of the differences in insect community composition are statistically significant. Again, a repeat of this experiment with larger sample sizes would help distinguish whether the findings here are a robust ecological pattern, or whether the differences in insect attraction between colored LEDs are merely due to random chance.

Our mushroom study species attracted a different insect assemblage than was found in previous studies with different mushrooms. In this study, the bioluminescent mushrooms primarily attracted Diptera species, with a few Hymenoptera and several Unknown species. After conducting a similar study in Brazil with the bioluminescent mushroom *Neonothopanus gardneri*, Oliveira *et al* 2015 found *N. gardneri* attracted mostly insects in the Diptera, followed by Hymenoptera, Coleoptera, and finally Hemiptera (Oliveira et al. 2015). In Sivinski 1981, the author found that luminous *Mycena* sp. primarily attracted Isopods, with the second highest abundance being insects in the Collembola, and the third category Diptera (Sivinski 1981). It will be informative to perform this type of experiment on more bioluminescent mushrooms of different lineages, in different forest types. Such studies will help determine if differences in insect assemblage are due to differences in insect abundance at the sites, or whether different mushroom species have evolved to attract different assemblages of insects.

Research from other teams lend credence to the hypothesis that mushroom bioluminescence attracts insects. In the bioluminescent mushroom *Mycena chlorophos*, the light is constrained to the spore-producing pileus and gills (Teranishi 2016), further evidence that the role of light is related to spore dispersal. We find the light is similarly distributed in *M. sypherii* (Figure 2C). Olveira *et al* 2015 found that *N. gardneri* utilized circadian rhythm to time its luminescence to only light up at night, suggesting that the role of light was not an accidental byproduct of metabolism (Oliveira et al. 2015).

However, several lines of evidence indicate that mushroom bioluminescence does not always serve to attract insect dispersers. The mycelia of some North American species of *Armillaria* produce light while the mushroom does not (Mihail 2015). For example, *Armillaria mellea* possesses the full enzymatic machinery to produce bioluminescence in the mycelium, but in the fruiting body, the synthesis of the luciferin precursor is blocked (Mihail 2015). This species may be undergoing selection to lose bioluminescence. Moreover, luminescence of *A. gallica* can be stimulated by mechanical disturbance, indicating light may play an entirely different role, perhaps related to cell repair. Future work should explore the potential ecological, or physiological, role of bioluminescent mycelia.

In conclusion, we found what may be a new bioluminescent mushroom species in the high-elevation oak forests of Costa Rica. While we predicted that green lights would attract different insect assemblages than other colors, the differences were not statistically significant. However, we did find evidence that the bioluminescent mushroom and green LEDs attract insects, particularly Dipteran flies that would be well suited to move mushroom spores further than wind dispersal. Our findings corroborate a few other similar studies in tropical regions that found mushroom nightlights may attract insects. Future work is needed to confirm whether different LED colors attract different insect assemblages, and to determine the importance of forest type in driving insect attraction to bioluminescent mushrooms.

## Acknowledgements

First and foremost, we wish to express sincere gratitude to the Organization for Tropical Studies for organizing and hosting the course Fungi and Fungi-like Organisms. Thank you to the OTS course instructors, Greg Mueller and Priscilla Chaverri for their stellar teaching and guidance. We thank John Taylor for the gift of polycarbonate and related advice. Thank you to Brian Joseph for letting CA Adams borrow special drill bits that don’t shatter plastic. Finally, a sincere thank you to the members of the Taylor and Bruns labs for valuable feedback on the manuscript.

## Notes

### Competing Interest Statement

The authors have declared no competing interest.

